# Robust encoding of acoustic identity in alpaca hums - a basis for individual recognition

**DOI:** 10.1101/2024.05.31.596793

**Authors:** Kaja Wierucka, Stephan T. Leu

## Abstract

Individual recognition is an important element of social interactions among animals. While the presence of individually distinct vocalisations (providing a basis for individual recognition) has been widely tested across species, information about which components of a call encode this information is lacking. We investigated whether female alpaca (*Vicugna pacos*) vocalisations, particularly their hums, encode information about individual identity and explored which parameters contribute to this encoding. We recorded vocalisations from 9 adult female alpaca and extracted both spectro-temporal features (frequencies and duration) and mel-frequency cepstral coefficients (MFCC). Random forest analyses revealed clear individual differences in both datasets, with the spectro-temporal features allowing for slightly more accurate classification than MFCC (71% and 66.5% accuracy for spectro-temporal features and MFCC, respectively). These robust acoustic identities have the potential to provide a basis for individual recognition in alpaca, which could have important flow on effects for alpaca communication, as it allows receivers to modulate their response to the caller’s identity. Alpaca, as herd-living and vocal animals, provide an excellent model system for better understanding the mechanisms, causes and consequences of recognition and inter-individual communication.

## INTRODUCTION

Recognition plays a fundamental role in animal communication systems. It influences the occurrence and nature of many social interactions among individuals, including mating, parental care, shared vigilance, and social foraging (Enquist et al., 2010). Recognition varies in specificity, with animals distinguishing broader categories such as species, sex or age, kin and familiar/non-familiar, or even specific individuals (Yorzinski, 2017). Individual recognition is the most complex and cognitively challenging form of recognition, as it requires individuals to identify others based on individually distinctive characteristics. This requires differentiating and remembering the cues or its elements that encode this information and accurately assigning it to a template representing an individual (Beecher, 1989; Tibbetts and Dale, 2007). Individual recognition is especially beneficial for group-living species, where repeated interactions among individuals are common and allows to modulate behaviours based on previous encounters and their outcomes (Yorzinski, 2017).

Individual identity information can be encoded within different sensory modalities (Bradbury and Vehrencamp, 2011) or multiple sensory channels at once (Higham and Hebets, 2013; Wierucka et al., 2018). Yet, it is acoustic cues that have been predominantly studied in the context of recognition, because they contain information about immediate states, can be used over short and long distances and tend to propagate through the environment better than visual or olfactory cues (Bradbury and Vehrencamp, 2011). Although bioassays, such as playbacks, are used to confirm that animals can identify conspecifics, they are often challenging to conduct. Therefore, as individually distinct characteristics in acoustic cues are necessary for recognition, an initial step is to analyse the structure of calls to identify any structural differences that could encode individual identity. While the encoding of individual identity information has been demonstrated in many mammalian taxa (Carlson et al., 2020; Linhart et al., 2022; Yorzinski, 2017) the specific components of a cue that encode individual identity and the robustness of these cues remain largely unknown.

The source and filter theory (Fant, 1970; Titze, 1994) links vocal production, acoustic structure and the perception of cues and thereby predicts which elements of a vocal cue are more likely to encode information. In mammals, the glottal wave is produced in the larynx (source) through the vibration of vocal folds and is then modified by the vocal tract (filter; Titze, 1994). Therefore, source-related features typically refer to measurements derived from the pitch, duration, tempo and amplitude contour, while the filter-related parameters include those related to the spectral envelope (e.g., formant frequencies, Mel-frequency cepstral coefficients; MFCC) of the acoustic signal (Taylor et al., 2016). Determining the origin of the identity cue for those taxa may shed light onto the function of specific vocalisations and aid in determining how and why given cues evolved. Nevertheless, many studies fail to integrate both source- and filter-related parameters in their analyses, neglecting to examine the relative significance of these factors in encoding identity (but see: Owren et al., 1997; Stoeger and Baotic, 2016; Vannoni and Mcelligott, 2007).

Source-related measurements are more flexible, and variable yet have been previously shown to play a role in individual, kin, group and mother-offspring recognition (reviewed in Taylor et al., 2016). Filter-related measurements have been shown to serve as an honest cue in some species, e.g., koalas (Charlton et al., 2011, 2012), domestic dogs (Taylor et al., 2010), pinnipeds (Sanvito et al., 2007), pandas (Charlton et al., 2010) primates (Fitch, 1997; Harris et al., 2006), deer (Reby et al., 2005; Vannoni and Mcelligott, 2008). However, as the filter-related acoustic characteristics of the produced sound reflect individual morphology and cannot be easily modified without a flexible vocal tract, they tend to be a more stable and reliable cue and are therefore more predictable (Taylor et al., 2016). This information can be used by animals directly for individual recognition or to make the recognition process more efficient by distinguishing broader categories of individuals first (e.g., size, sex, age) and then identifying the individual. Parameters related to the spectral envelope (e.g., MFCCs) have been extensively used for voice recognition in humans and have been shown to be an important cue for recognition in primates (Clink et al., 2019; Clink and Klinck, 2021; Gamba et al., 2012; Mielke and Zuberbühler, 2013) as well as other taxa (Townsend et al., 2014).

Group-living species make excellent communication study models, as more complex social interactions (usually requiring recognition) result in more elaborate communication (Freeberg et al., 2012; Peckre et al., 2019). However, studying communication in social animals can often be challenging due to the technical and practical difficulties of obtaining repeated, clean recordings from multiple individuals, produced in a natural behavioural context. Captive animals may exhibit behaviours that may be altered due to their repeated exposure to humans. However, obtaining high quality recordings is more feasible as animals are easily accessed and movement restricted to an enclosure/cage. Domesticated herd mammals that can freely move in large paddocks, present a unique opportunity to obtain a dataset that is relatively high in sample size, and provides clear recordings with behaviours being unaffected by data collection with the use of animal-mounted tags. Alpaca (*Vicugna pacos*), allowing for collection of acoustic data in natural behavioural contexts are an excellent study species for investigating animal communication that provides unparalleled opportunities for obtaining insights into the mechanisms of recognition and communication.

Alpaca are herd-living camelids from South America, that form stable social groups, with members engaging in repeated interactions (Miranda-de la Lama and Villarroel, 2023). They have been domesticated and are farmed primarily for the purpose of fibre production, gaining popularity particularly among smaller farms and private owners (Alonso, 2010). They are more social than llamas (*Lama glama*) and are known to produce a variety of vocalisations including hums, snorts, screams, screeches and grunts (Fowler, 2013). The hum is the predominant call type, particularly between group members and serves various functions (Fowler, 2013), including as contact calls in the species (Kapustka et al., 2018). In this study, we aimed to determine whether alpaca hums have the capacity to encode information about individual identity and whether identity cues were primarily encoded in spectro-temporal features (representing primarily source-related features, or MFCC (representing primarily filer-related parameters).

## METHODS

### Study site

We conducted our study at the livestock facilities of the University of Adelaide (Roseworthy campus), Australia. The fenced study paddock was approximately one hectare (140 m x 70 m) in size. Throughout the study, animals had access to shaded areas, hay and fresh water. No other animals occupied the paddock during the study.

### Recording alpaca vocalisations

Our study herd consisted of 12 adult female alpaca, of which 9 were used in the analyses (Table 1). All animals were familiar with each other as they were kept together as a herd. The genetic relatedness was unknown; parts of the herd originated from different breeders. All animals were accustomed to human handling. In December 2020, we equipped each of the alpacas with a small audio recorder (model: VIM-PWBVOICE2 by VIMEL Security Solutions, recorded at 48kHz) and Global positioning system (GPS; i-gotU GT-120 by MobileAction, with a larger battery CE04381 by Core Electronics). The head halter, including the devices, weighed a mean of 303g (SE 2.5g, n=12). The audio recorder and its microphone were mounted close to the mouth of the individual, assuring a high amplitude of the vocalisations of the focal animal compared to other individuals. Audio was continuously recorded for approximately 6 hours each day, for 8 days. In the evening, we removed all halters to allow for the downloading of the data and recharging of the battery. All animals were re-haltered in the morning. Individual alpaca could be identified by their already present unique ear tag number as well as distinct pelt colouration. At the end of the study, we removed all halters and released the alpaca back into their paddock. All animals were treated using procedures formally approved by The University of Adelaide Animal Ethics Committee (approval number S-2020-044).

**Table 1.**
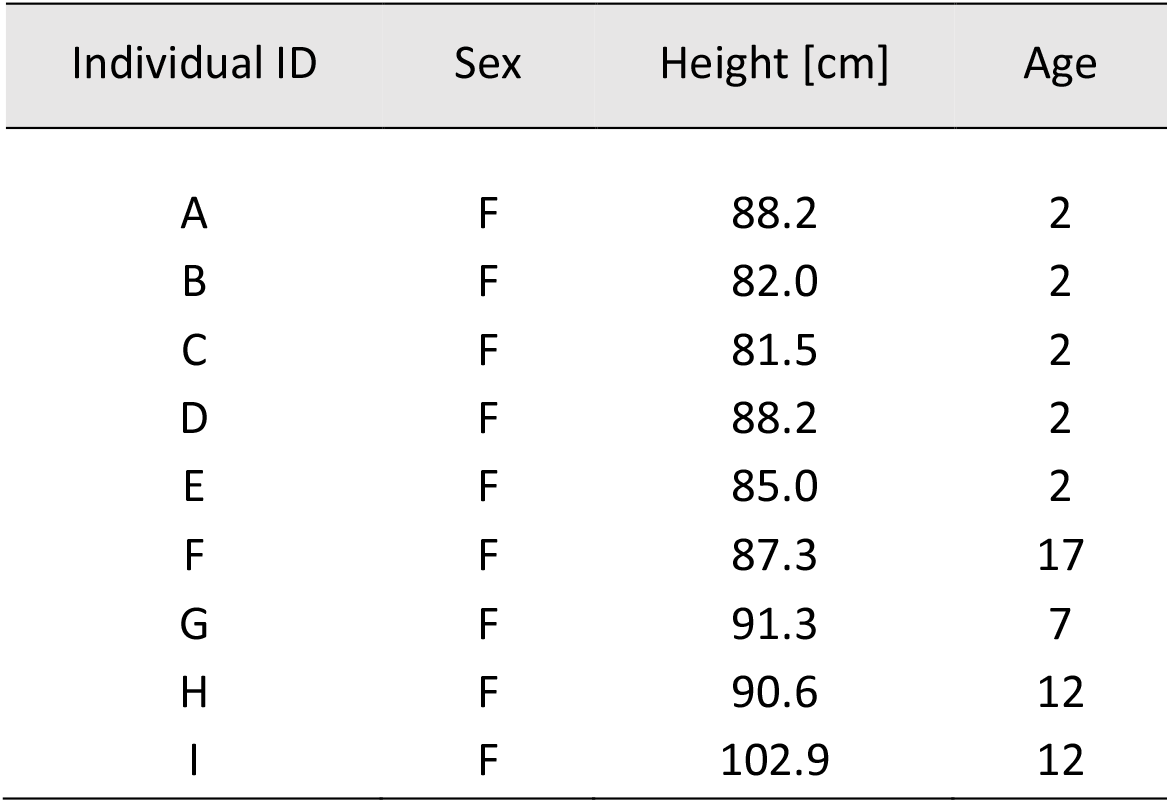
Individual traits of the 9 study subjects. F – female; Height – cm at withers; Age - years.

### Acoustic processing

All recordings were processed in RavenPro 1.6. Spectrogram settings were set to a Hann window of 41.7 ms, with a hop size of 20.8 ms and an overlap of 50% (DFT size 2048 samples). We took a semi-automated approach, where calls were segmented manually, but measurements were extracted automatically with RavenPro 1.6. We extracted all calls that had a predominantly flat peak frequency contour, which we refer to as hums (some included a frequency modulated fragment at the beginning and/or end of the call; Figure 1). Hums are the most frequently produced call type by alpaca, used in social contexts (Fowler, 2013). We limited the dataset to high quality calls only, clearly visible on the spectrogram, with no overlap or background noise. We excluded three individuals from the analysis for which we obtained less than 10 high quality vocalisations. The remaining individuals differed in the number of high-quality vocalisations, ranging from 14 to 134.

**Figure 1.**
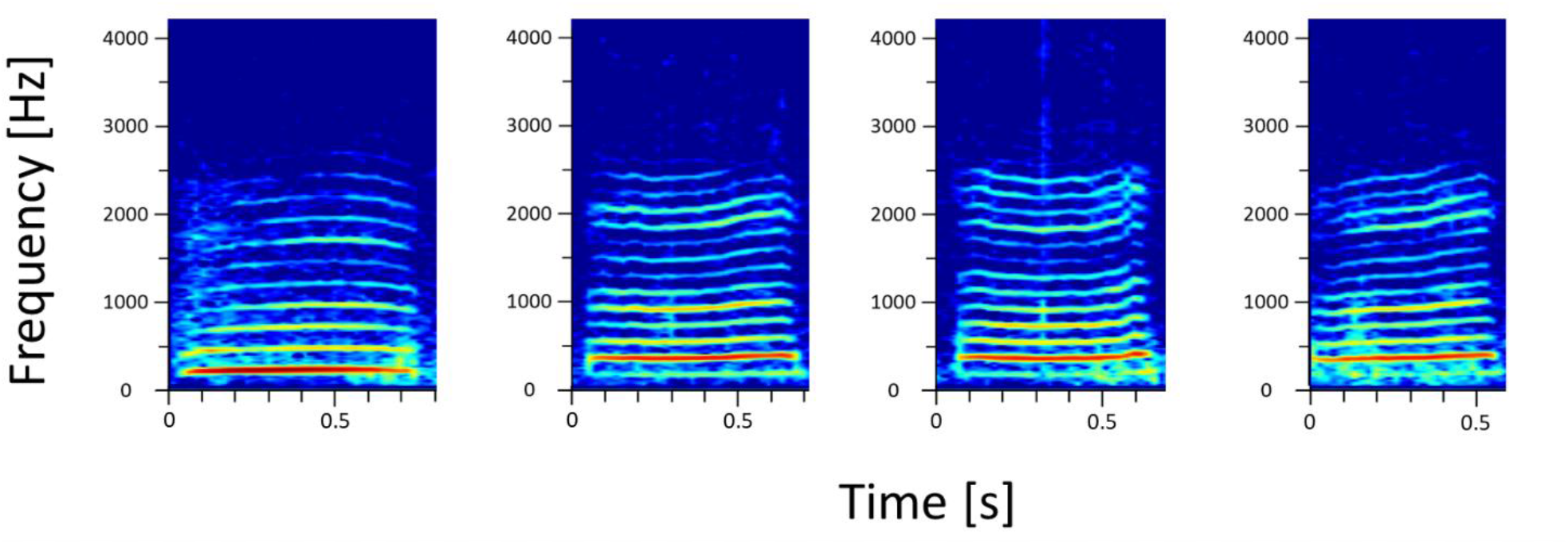
Examples of the diversity of alpaca vocalisations classified as hums. Spectrogram settings: DFT size 2048 samples, Hann window of 41.7 ms, with a hop size of 20.8 ms and an overlap of 50%.

To ensure consistency across feature extraction, measures were extracted automatically from the vocalisations. We extracted both frequency parameters and MFCC from bandstop filtered calls (filter 0-220Hz, to remove low frequency noise. All frequencies of hums occur above 220Hz). The spectro-temporal parameters convey mostly information related to the fundamental frequency (F0) and therefore the source-related features of the vocalisation. MFCCs are measures of the spectral envelope of the call. These, in part, correspond to the filter-related features of the vocalisation (e.g., peaks in the spectral envelope are formant frequencies; Taylor et al., 2016), and have been shown to have the potential for encoding individual identity in other herd animals (e.g., goats (Briefer and McElligott, 2011), sheep (Wierucka et al., 2024)). To avoid bias resulting from observer selections and consequent inconsistencies in spectral frequency measurements, we extracted robust measurements in RavenPro 1.6: frequency 5%, frequency 95%, and bandwidth 90% and duration 90% (defined in Charif et al., 2010; following Wierucka et al., 2021). These measurements take the point of measurement based on the energy that is stored in the call rather than based on box selection boundaries, making the measurements more robust. In addition to these measurements, we also included the peak frequency, average entropy (extracted in RavenPro 1.6) as well as the F0 (‘lowest dominant frequency band’ extracted in package soundgen; Anikin, 2019). MFCC were extracted in R, using the BehaviouR package (Clink, 2020). Twelve cepstra were calculated per window (corresponding to the spectral envelope, rather than higher coefficients that relate to the glottal pulse of the sound) within the 0-5000Hz range (as hums were contained within it).

### Statistical analysis

To assess the encoding of individuality, we implemented a random forest classifier separately for the spectro-temporal and MFCC datasets. Data were split 70:30 into training and testing sets (within individual and with no overlap), with results reported for testing sets only. We ran a random forest model (randomForest::tuneRF; Liaw and Wiener, 2002) using the optimal mtry found (doBest=TRUE), with 1000 trees used at the tuning step. Based on the confusion matrices obtained from the classifiers, we calculated overall accuracy and by-class (by-individual) F1 scores using the caret package (Kuhn and Max, 2008). We evaluated the importance of each variable (only for the spectro-temporal dataset, as MFCC variables are difficult to interpret in a biologically meaningful way on their own) for the classification task based on the Mean Decrease Accuracy values. These values determine how much the prediction error would increase if that variable was changed (Liaw and Wiener, 2002).

All animals were adult females, yet they varied slightly in body size. Hence, we tested whether body size differences could have contributed to any differences in acoustic cue structure among individuals. We related animal body size (height at withers in centimetres) with the average F0 measurement of hums produced by each individual using a Spearman’s rank correlation (Bowling et al., 2017). MFCC are less easily explained in terms of biological relevance, therefore, to establish whether any of the variables were correlated with body size, we reduced the number of variables with a principal component analysis (PCA). We retained PCs with eigenvalues over 1 and calculated the mean values of those PCs for each individual. We then related these mean values with individual body size using a Spearman’s rank correlation and adjusted the p values for multiple comparisons (fdr adjustment method).

All statistical analyses were conducted in R version 4.0.1 (R Core Team, 2020).

## RESULTS

Vocalisations had an average F0 (±SE) of 272±7.5 Hz, duration 90% of 464 ±0.06 ms and peak frequency of 353 ±22 Hz (Table 2). All raw data extracted and used for analysis are provided in Supplementary Materials.

**Table 2.** Mean acoustic spectral features across hums recorded for each individual.

| Individual | Fundamental Frequency (Hz) | Duration 90% (ms) | Peak Frequency (Hz) | Frequency 5% (Hz) | Frequency 95% (Hz) | Bandwidth 90% (Hz) |
| --- | --- | --- | --- | --- | --- | --- |
| A | 284 | 380 | 403 | 259 | 705 | 445 |
| B | 279 | 366 | 308 | 244 | 795 | 551 |
| C | 305 | 390 | 363 | 301 | 613 | 311 |
| D | 290 | 362 | 402 | 265 | 700 | 435 |
| E | 244 | 584 | 331 | 224 | 926 | 701 |
| F | 318 | 569 | 382 | 286 | 1164 | 877 |
| G | 229 | 687 | 271 | 209 | 959 | 750 |
| H | 220 | 375 | 412 | 213 | 807 | 594 |
| I | 277 | 463 | 295 | 259 | 787 | 527 |
| overall mean | 272 | 464 | 352 | 251 | 828 | 577 |

The random forest classified hums to individual alpaca identity based on the spectro-temporal features with an accuracy of 71% (by-class F1 scores ranging 0.417-0. 882; Supplementary Materials Table 1). The classification accuracy based on MFCC values was 66.5% (by-class F1 scores ranging 0.363-0.829; Supplementary Materials Table 1). F0 was the most important variable for the classifier using the spectro-temporal parameters of calls (Supplementary Materials; Table 2). Finally, we observed no significant correlation of body size and F0 (p=0.1447). The PCA identified 30 PCs, none of which were significantly correlated with height (Supplementary Materials; Table 3).

## DISCUSSION

Individual recognition is an essential element of animal sociality (Tibbetts and Dale, 2007). However, how individuality is encoded can vary by species, call type as well as context. Alpaca are a group-living social mammal and we investigated whether the hum, a commonly produced call type in alpaca, encodes individual identity information. We often lack information about the importance of various call components in conveying this information. Here, we have shown that in the alpaca hum, individuality is encoded in both spectro-temporal features of the call as well as MFCCs (describing the spectral envelope of the sound). The classifier based on spectro-temporal features was slightly more accurate, with the F0 being the most important feature for determining individual distinctiveness. Importantly the acoustic individual differences did not reflect body size differences. This suggests that alpaca hums they have the capacity to encode true individual identity rather than simply size differences.

Our models were able to classify the alpaca hums to individuals based on spectro-temporal parameters as well as MFCC values with higher accuracy than expected by chance. As MFCC have been developed for the study of human language, in the past, parameters related to the spectral envelope of a wave have been primarily used for studying individual identity encoding in primates (e.g., Clink et al., 2020, 2021; Coye et al., 2022; Fedurek et al., 2016; Mielke and Zuberbühler, 2013). Now, filter-related measures have become an increasingly popular metric for determining individual identity encoding in calls of other mammals, e.g., bats (Prat et al., 2016), elephants (Stoeger and Baotic, 2016), mongooses (Rubow et al., 2017), and guinea pigs (Verzola-Olivio et al., 2021). However, few studies incorporate both source- and filter-related parameters into their analyses to look at the relative importance of these for encoding identity (but see: Owren et al., 1997; Stoeger and Baotic, 2016; Vannoni and Mcelligott, 2007).

Encoding information in multiple characteristics of a cue contributes to its robustness and ensures the transfer of information when other factors affect the produced call (Ay et al., 2007; Bro-Jørgensen, 2009; Hebets and Papaj, 2005). For example, while call pitch commonly serves as a distinguishing trait for individual identity (e.g., Blackshaw et al., 1996; Gros-Louis, 2006; Palacios et al., 2007; Wierucka et al., 2021; Wijers et al., 2021), fear-induced heightened arousal can lead to pitch alterations (Taylor et al., 2016). While the main purpose of an alarm call is typically to signal immediate threat, animals could benefit from possessing knowledge of the caller’s identity, thereby enabling assessments of signal reliability (Blumstein and Daniel, 2004). The alpaca hum is important in a social context, and has been suggested to be a contact call, but also that it is used during separation (Fowler, 2013). Contact calls typically encode individual attributes (Kondo and Watanabe, 2009), which is consistent with our finding that the hums differed between individuals and encoded individual identify. Additionally, our investigation reveals substantial individual distinctiveness within alpaca hums, even when applying a broadly defined hum category and acknowledging some degree of structural variability in the calls (Figure 1). This observation underscores the robust nature of identity encoding within this call type.

Our findings demonstrate that in alpaca both MFCC and spectro-temporal features carry identity information and could be used for individual recognition. MFCC are known to serve the function of honest cues of body size in mammals (Taylor and Reby, 2010), yet our correlation tests confirmed that body size did not drive the observed individual differences for neither source-nor filer-related features. We found a more pronounced contribution of individual identity in differences found in spectral features, which could point to their higher importance in individual recognition in alpaca, and perhaps a more common usage of these elements of the cue. This is consistent with other mammal species, where spectral frequency has been shown to play an important role in individual distinctiveness of cues or recognition (e.g., Blackshaw et al., 1996; Gros-Louis, 2006; Palacios et al., 2007; Wierucka et al., 2021; Wijers et al., 2021). The fundamental frequency contributed most to the observed differences between alpaca, which is consistent with previous findings in e.g., wolves (Palacios et al., 2007) and hyenas (Mathevon et al., 2010). However, we also found that individuality is encoded in MFCCs of alpaca hums, indicating that the spectral envelope of the sound (and potentially filter-related parameters) of the cue also play an important role in individual recognition and are not just used for broader category assessment.

Classification success varied among individuals, which we can likely explain by natural variation in the distinctiveness of vocalisations. However, we cannot exclude the possibility that this variation may also result of vocal similarities among some individuals. Vocal similarity may be a result of many factors, including social or environmental accommodation (Ruch et al., 2018), genetic relatedness (Hauser, 1996), or population processes such as turnover rates (Parker et al., 2022). It is possible that certain individuals in our study associate with each other more frequently and therefore their calls converge (as summarised in Ruch et al., 2018), or that a more similar genetic make-up results in similar build and therefore sounds produced (e.g., in pandas (Charlton et al., 2009) or mouse lemurs (Zimmermann and Hafen, 2001)). Investigating the correlations between social interaction patterns, relatedness and vocal signatures is an interesting avenue for future studies that would help shed light onto the characteristics of the vocal communication network and aid in linking two essential elements of within-group dynamics – social propensity and communication.

Our findings point to analytically detectable individual differences in alpaca hums. While this suggests that the vocalisations convey information on individual identity during alpaca communication, bioassays, such as playback experiments, would confirm that alpaca are able to detect those differences and respond to them. Interestingly, hum fundamental frequency was 271Hz on average. While this is within the hearing range of alpaca, their highest acoustic sensitivity is at 8kHz (Heffner et al., 2014), pointing to potentially other acoustic cues or elements of higher frequency within cues being of importance for communication.

Alpaca form stable social groups and interact repeatedly with group members (Miranda-de la Lama and Villarroel, 2023). In complex social groups, including in alpaca, not all individuals have the same influence on the group. Some individuals have a disproportionately large influence and occupy leadership positions in the group. Similarly, alpaca establish a social hierarchy within the group, with high- and low-ranking individuals (Miranda-de la Lama and Villarroel, 2023). Being able to distinguish between individuals, such as our auditory cues, would allow animals to modulate their social interactions based on past experiences with those individuals. For instance, individuality in alarm calls would allow to modulate the response depending on the reliability of the caller. The alpaca hum is an auditory contact among group members, likely operating at relatively shorter distances given its features, than alarm calls. Hence, for instance it may facilitate mother-offspring recognition, or group coordination through being able to distinguish between leader and follower individuals. However, the function of the individuality of the hums still needs to be established. Nevertheless, for individual identity in calls to evolve the response by the receiver needs to be beneficial for the sender, otherwise selection would favour concealing identity (Sheehan et al., 2014).

Our findings provide an important foundation to understanding alpaca auditory communication. They add to a growing awareness of information on individual identify, here based on robust auditory cues. Further research on how alpaca use this information to modulate dyadic social interactions and how this may upscale to the social network structure would provide important and interesting new insights into complex animal societies, and how they are maintained.

## ACKNOWLEDGEMENTS

This work was supported by a School of Animal and Veterinary Sciences seed grant to STL. We would like to thank Cameron McKinnon for his assistance with building the halters and collaring the alpaca, and SAVS staff for animal husbandry.

## ETHICS APPROVAL

All animals were treated using procedures formally approved by The University of Adelaide Animal Ethics Committee (approval number S-2020-044).

