## Supplementary material for "Robust encoding of acoustic identity in alpaca hums - a basis for individual recognition": Supplemenary Materials

Table 1. By-class F1 scores calculated for a classification task using a random forest algorithms to determine the accuracy of classification of alpaca vocalisations to individual identity. Two models were run, one using spectro-temporal features and one using MFCC values.

| Features used | Individual | | | | | | | | |
|---|---|---|---|---|---|---|---|---|---|
| | A | B | C | D | E | F | G | H | I |
| Spectro-temporal | 0.714 | 0.467 | 0.847 | 0.417 | 0.581 | 0.706 | 0.742 | 0.882 | 0.478 |
| MFCC | 0.667 | 0.462 | 0.763 | 0.364 | 0.421 | 0.571 | 0.667 | 0.829 | 0.467 |

Table 2. Variable importance for a classification task using a Random Forest algorithm. Spectro-temporal parameters were used.

| Variable | Mean Decrease Accuracy |
|---|---|
| Fundamental Frequency (Hz) | 44.573 |
| Frequency 5% (Hz) | 23.418 |
| Peak Frequency (Hz) | 19.635 |
| Average entropy (bits) | 16.513 |
| Frequency 95% (Hz) | 7.286 |
| Bandwidth 90% (Hz) | 11.066 |
| Duration 90% (ms) | 5.919 |

Table 3. Relationship between the acoustic structure of alpaca hums and individual height. Results show the values of Spearman's correlation tests for each PC with p-values adjusted for multiple comparisons. PCs were obtained following applying a PCA on MFCC extracted from each call (we retained PCs with eigenvalues over 1 and calculated the mean values of those PCs for each individual; see methods for details).

| PC | rho | p value |
|---|---|---|
| PC1 | 0.192 | 0.620 |
| PC2 | -0.460 | 0.213 |
| PC3 | -0.494 | 0.177 |
| PC4 | -0.335 | 0.379 |
| PC5 | 0.310 | 0.417 |
| PC6 | -0.067 | 0.864 |
| PC7 | 0.661 | 0.053 |
| PC8 | -0.176 | 0.651 |
| PC9 | 0.393 | 0.295 |
| PC10 | -0.285 | 0.458 |
| PC11 | 0.226 | 0.559 |
| PC12 | -0.042 | 0.915 |
| PC13 | 0.368 | 0.330 |
| PC14 | 0.084 | 0.831 |
| PC15 | 0.661 | 0.053 |
| PC16 | -0.544 | 0.130 |
| PC17 | -0.711 | 0.032 |
| PC18 | -0.410 | 0.273 |
| PC19 | -0.310 | 0.417 |
| PC20 | -0.519 | 0.152 |
| PC21 | -0.444 | 0.232 |
| PC22 | 0.050 | 0.898 |
| PC23 | 0.377 | 0.318 |
| PC24 | -0.067 | 0.864 |
| PC25 | 0.176 | 0.651 |
| PC26 | -0.192 | 0.620 |
| PC27 | 0.301 | 0.431 |
| PC28 | -0.142 | 0.715 |
| PC29 | 0.042 | 0.915 |
| PC30 | 0.318 | 0.404 |
